# A natural bacterial pathogen of *C. elegans* uses a small RNA to induce transgenerational inheritance of learned avoidance

**DOI:** 10.1101/2023.07.20.549962

**Authors:** Titas Sengupta, Jonathan St. Ange, Rebecca Moore, Rachel Kaletsky, Jacob Marogi, Cameron Myhrvold, Zemer Gitai, Coleen T. Murphy

**Affiliations:** Lewis Sigler Institute for Integrative Genomics, Princeton University, Princeton, NJ 08544, USA; Department of Molecular Biology, Princeton University, Princeton, NJ 08544, USA

## Abstract

Previously, we discovered that a small RNA from a clinical isolate of *Pseudomonas aeruginosa*, PA14, induces learned avoidance and its transgenerational inheritance in *C. elegans. Pseudomonas aeruginosa* is an important human pathogen, and there are other *Pseudomonads* in *C. elegans’* natural habitat, but it is unclear whether *C. elegans* ever encounters PA14-like bacteria in the wild. Thus, it is not known if small RNAs from bacteria found in *C. elegans’* natural habitat can also regulate host behavior and produce heritable behavioral effects. Here we found that a pathogenic *Pseudomonas vranovensis* strain isolated from the *C. elegans* microbiota, GRb0427, like PA14, regulates worm behavior: worms learn to avoid this pathogenic bacterium following exposure to GRb0427, and this learned avoidance is inherited for four generations. The learned response is entirely mediated by bacterially-produced small RNAs, which induce avoidance and transgenerational inheritance, providing further support that such mechanisms of learning and inheritance exist in the wild. Using bacterial small RNA sequencing, we identified Pv1, a small RNA from GRb0427, that matches the sequence of *C. elegans maco-1*. We find that Pv1 is both necessary and sufficient to induce learned avoidance of Grb0427. However, Pv1 also results in avoidance of a beneficial microbiome strain, *P. mendocina*; this potentially maladaptive response may favor reversal of the transgenerational memory after a few generations. Our findings suggest that bacterial small RNA-mediated regulation of host behavior and its transgenerational inheritance are functional in *C. elegans’* natural environment, and that different bacterial small RNA-mediated regulation systems evolved independently but define shared molecular features of bacterial small RNAs that produce transgenerationally-inherited effects.

## Introduction

Plants and animals have evolved diverse mechanisms to adapt to constantly changing environmental stimuli. Some of these stimuli are encoded as molecular changes that do not involve changes in DNA sequence, but are instead epigenetic, that is, mediated through changes in non-coding RNAs, DNA modifications, histone modifications, and nucleosome positioning (1–6). These changes can occasionally cross the germline and confer adaptive benefits to the first generation of progeny (intergenerational) (7–20) or more (transgenerational) (21–33). Multigenerationally-inherited effects can provide adaptive advantages in changing environments, particularly in organisms with short generation times (34–36).

Over the past decade, instances of multi-generational inheritance have been reported in various organisms (37–39). We previously characterized an example of epigenetic inheritance in response to a physiological stimulus, highlighting its adaptive benefits in *C. elegans*: upon exposure to the pathogenic *Pseudomonas aeruginosa* strain PA14, worms learn to subsequently avoid the bacteria, then pass on this learned avoidance to four generations of progeny (27). A single small RNA from PA14, P11, mediates this avoidance and its transgenerational inheritance through reduction of the worm neuronal gene *maco-1*, which results in a switch from attraction to avoidance behavior (23,26). These studies provided the first example of bacterial small RNA-mediated regulation of a learned behavior and its transgenerational inheritance (23,26,27). However, PA14 is a human clinical *Pseudomonas* isolate; whether bacteria in *C. elegans’* natural environment elicit learned responses and multi-generational inheritance of learned responses through small RNAs is not known.

Diverse bacterial species influence *C. elegans* physiology and life history traits (40–42). Bacterial species from *C. elegans*’ natural environment have been systematically characterized (43–49). These studies revealed multiple features of the microbiota in *C. elegans* natural habitat and their relationship to host physiology (50,51). Studying bacterial species that are naturally associated with *C. elegans* might reveal processes that occur in the wild, and not in the laboratory, and vice versa; for example, bacteria from *C. elegans’* natural environment suppress mortal germline phenotypes that wild worms exhibit on laboratory strains of *E. coli* (52). Therefore, it is important to test the physiological relevance of laboratory experimental results under more natural conditions.

Bacterial species in the worm microbiome that induce stress and immune response reporters are categorized as pathogenic (47), while species that promote increased worm growth rates are categorized as beneficial (47,49,53), and some of these species confer protection against pathogenic species (44,54). Other species are beneficial in some contexts and pathogenic in others (55). Therefore, it may be evolutionarily favorable for worms to have plastic responses to different bacterial classes that they naturally encounter. Worms are naively attracted to specific beneficial and neutral bacterial species (e.g., *Pseudo-monas mendocina* and *Proteus mirabilis*, respectively) when given a choice between these bacteria and their laboratory diet *E. coli* HB101(56). Similarly, worms grown on the beneficial bacterial strain *Providentia alcalifaciens* prefer this bacterial species over their laboratory diet *E. coli* OP50 in a behavioral choice assay (57). These beneficial bacterial species modulate *C. elegans’* attraction towards several chemicals. Naïve or learned attraction in response to beneficial bacteria that worms encounter in their natural environment may have evolved as an evolutionarily favorable strategy. However, it is not known if worms can learn to avoid the various pathogenic bacterial species in their environment, or inherit this learned avoidance. Additionally, whether bacteria in *C. elegans’* natural environment can modulate the host nervous system through small RNAs and whether they can induce transgenerationally-inherited effects are not known.

*Pseudomonas* is one of the largest among the bacterial genera that constitute *C. elegans’* natural microbiome (47). In this study, we examined *C. elegans’* behavioral responses to a Pseudomonad species present in its natural microbiome. We found that an isolate of *Pseudomonas vranovensis* can elicit learned avoidance and its transgenerational inheritance through a single small RNA that is both necessary and sufficient. However, this learned response to *P. vranovensis* also leads to avoidance of a beneficial bacteria also found in *C. elegans’* environment, *P. mendocina*. Our work reveals a transgenerational effect in response to bacteria in *C. elegans’* natural microbiome, underscoring the physiological relevance of transgenerational inheritance and its significance in the wild. We also identified a new small RNA that can induce a learned behavior in *C. elegans*, therefore expanding the repertoire of bacterial small RNA-mediated regulation of the host nervous system and helping to identify characteristics of small RNAs necessary for trans-kingdom signaling. Finally, the induced avoidance of a beneficial bacteria after pathogen training suggests that “forgetting” learned pathogen avoidance after a few generations might benefit *C. elegans*, limiting maladaptive behaviors.

## Results

### Wild microbiome bacteria induce learned avoidance

To examine if exposure to bacteria from *C. elegans’* natural environment can produce stereotypic behavioral responses and further, whether these could potentially be small RNA-mediated, we were curious about *C. elegans’* response to strains that are present in its natural microbiome (47). We chose nine different bacterial species, mostly from the CeMBio collection (50) to test. These include bacteria that (i) are pathogenic or impair worm growth and development (GRb0427, Jub19, Bigb0170), (ii) are beneficial, as they enhance worm growth rates or provide protection against pathogen infection (MSPm1, Cen2ent1, Jub38), or (iii) have positive or neutral effects depending on the physiological context (Myb71, Myb10, Jub66) (46,47,47,49,50,50). Starting as late L4 animals, we exposed *C. elegans* for 24hrs either to OP50 *E. coli* (the standard lab cultivation strain) or to the test bacteria, and then assayed their preference to OP50 vs. the test strain (**Figure 1A**). Prior to exposure to this pathogen, in general, *C. elegans* prefer the wild strains - both beneficial and pathogenic - relative to the laboratory food *E. coli* OP50 (OP50-trained, **Figure 1A-J**). After training, six of the bacteria showed no significant change in preference, despite the fact that two of those strains, Jub19 and Bigb0170, have detrimental effects on worms (47,50) ; however, 24hr of exposure to the remaining three strains (GRb0427, Myb71, and Cen2ent1) induced significant avoidance in the trained mothers (P0) (**Figure 1D, E, G**). That is, upon cultivation on a bacterial lawn for 24 hours, worms learn to robustly avoid the bacteria, as shown in a choice assay between OP50 and the test strain. Thus, it seems that most wild strains are inherently attractive to *C. elegans*, whether they are beneficial, neutral or pathogenic, and only a subset of strains induce avoidance in the P0.

**Figure 1.**
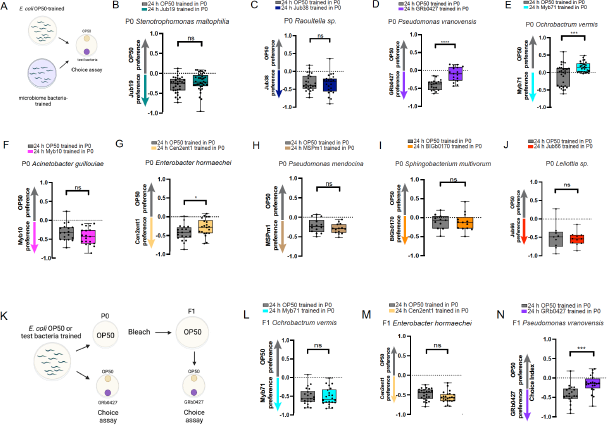
Wild microbiome bacteria induce learned avoidance. **(A)** Worms trained for 24 hours on *E. coli* OP50 or a test wild microbiome bacterial strain are tested in a choice assay between OP50 and the test bacterial strain. **(B-J)** Choice assays before and after training on *Stenotrophomonas maltophilia* Jub19 (B), *Raoultella sp*. Jub38 (C), *Pseudomonas vranovensis* GRb0427 (D), *Ochrobactrum vermis* Myb71 (E), *Acinetobacter guillouiae* Myb10 (F), *Enterobacter hormaechei* Cen2ent1 (G), *Pseudomonas mendocina* MSPm1 (H), *Sphingobacterium multivorum* Bigb0170 (I), *Leliottia sp*. Jub66 (J). **(K)** Worms trained for 24 hours on *E. coli* OP50 or a microbiome bacterial strain are bleached to obtain eggs, which are allowed to grow to Day 1 adults on OP50 plates. These adult F1 progeny are tested in a choice assay between OP50 and the respective bacterial strain. **(L-N)** Choice assays with F1 progeny of OP50 and Myb71-trained (L), OP50 and Cen2ent1-trained (M), and OP50 and GRb0427-trained (N) animals. Each dot represents an individual choice assay plate. Boxplots: center line, median; box range, 25th–75th percentiles; whiskers denote minimum-maximum values. Unpaired, two-tailed Student’s t test, ^****^p < 0.0001, ***p < 0.001, and *p<0.05, ns, not significant.

To determine whether this learned avoidance is inherited by the next generation, trained mothers (P0) were bleached and their progeny (F1) were raised on OP50 *E. coli* until adulthood, then were tested for their choice (with no training) (**Figure 1K**). We observed that although neither Myb71 nor Cent1 progeny inherited the learned avoidance from their mothers (**Figure 1L, M**), progeny of GRb04270-trained mothers also avoided GRb0427 (**Figure 1N**). Of the various bacteria we tested, only the pathogenic *P. vranovensis* GRb0427 both learned avoidance and inherited the mother’s learned avoidance.

### *C. elegans* learn to avoid the natural bacterial pathogen, *P. vranovensis*

Despite worms’ naïve attraction to *Pseudomonas vranovensis* GRb0427, this bacterium is pathogenic to *C. elegans*: adult exposure to GRb0427 causes severe illness (**Figure 2A**) and significantly reduces survival to less than 2-3 days (**Figure 2B**), in contrast to *C. elegans’* normal lifespan of 2-3 weeks.

**Figure 2.**
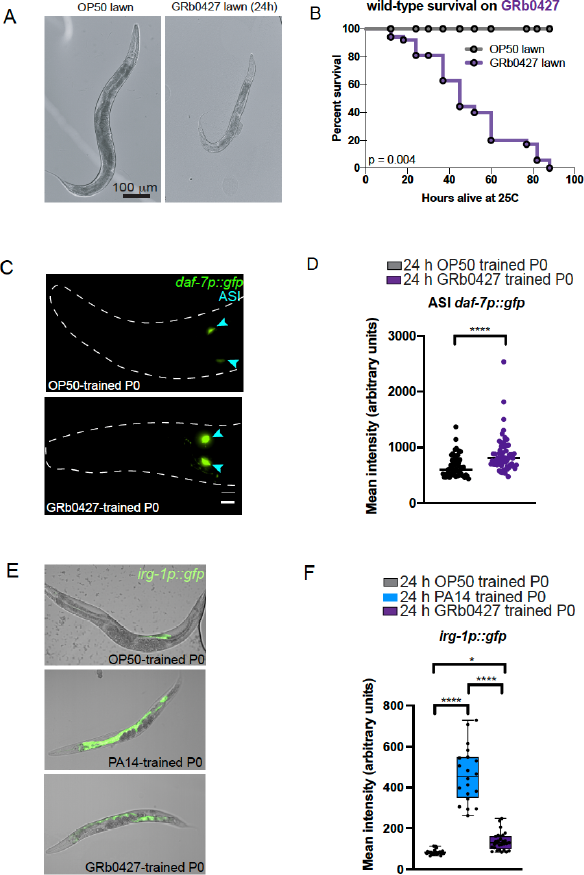
GRb0427, a natural Pseudomonad pathogen of *C. elegans* induces learned avoidance. **(A)** Representative images acquired after exposing Day 1 worms to OP50 (left) and GRb0427 (right) for 24 hours. GRb0427 is pathogenic and 24-hour exposure makes worms sick **(B)** Worms have significantly lower survival on a GRb0427 lawn compared to an OP50 lawn. **(C)** Representative image of a 24-hour OP50-trained or GRb0427-trained worm *expressing daf-7::GFP. daf-7::GFP* is expressed in the ASI sensory neurons. (blue arrowheads). The dashed line indicates the outline of the worm head. Scale bar = 10 μm. **(D)** Quantification of mean ASI *daf-7p::gfp* intensities from OP50 and GRb0427 animals shows higher expression in GRb0427-trained animals. **(E)** Expression of an *irg-1p::gfp* reporter in representative OP50-trained, PA14-trained and GRb0427-trained animals. Images are merged confocal micrographs of brightfield and GFP channels. Scale bar = 40 μm. **(F)** Mean fluorescence intensity of an *irg-1p::gfp* innate immune response reporter in OP50 (gray), *Pseudomonas aeruginosa* PA14 (blue), and GRb0427 (purple)-trained worms. Mean *irg-1::gfp* reporter intensity is significantly lower in GRb0427-trained worms compared to PA14-trained worms. Each dot represents an individual neuron (D) or an individual worm (F). Boxplots: center line, median; box range, 25th–75th percentiles; whiskers denote minimum-maximum values. Unpaired, two-tailed Student’s t test, ^****^p < 0.0001 (D); one-way ANOVA with Tukey’s multiple comparison’s test, ****p<0.0001, *p < 0.05 (F). For the survival assay in (B), ****p<0.001 (by Log-rank (Mantel-Cox) test for survival).

Exposure to *P. aeruginosa* PA14 causes gene expression changes in specific *C. elegans* sensory neurons; specifically, a 24-hour exposure to PA14 results in the induction of *daf-7p::gfp* expression in the ASJ neurons and an increase in *daf-7p::gfp* expression in the ASI neurons (27,58). PA14 small RNAs induce expression of *daf-7p::gfp* in the ASI that persists in the F1-F4 progeny generations (23,27), while the increase in ASJ *daf-7* is caused by PA14 secondary metabolites phenazine-1-carboxamide and pyochelin (58), and does not persist beyond the P0 (27). To determine whether altered *daf-7* levels correlate with the learned avoidance response to *P. vranovensis*, we examined *daf-7p::gfp* expression in GRb0427-trained animals (P0). Upon GRb0427 exposure, *daf-7p::gfp* levels significantly increase in the ASI neurons (**Figure 2C-D)**, but no expression was observed in the ASJ neurons, in contrast to the response to PA14 training (27) (**Figure S1**). Unlike PA14 and other pathogenic bacteria, exposure to GRb0427 triggers a significantly milder innate immune response, as indicated by low expression of a reporter of the *irg-1* gene reporter (**Figures 2E-F**). Lack of induction of phenazine-mediated ASJ *daf-7p::gfp* expression and only a mild induction of the *irg-1* dependent innate immune pathway suggest that these innate immune pathways might not play a significant role in the neuronal response to the wild bacteria GRb0427, even in the P0 generation, unlike the response to the clinical isolate PA14.

### The avoidance response to *P. vranovensis* is transmitted for four generations

The learned avoidance induced by a PA14 lawn and the PA14 small RNA P11 are communicated to naïve F1 progeny (27), and *daf-7* levels are also increased in the ASI neurons in the parental and four subsequent (F1-F4) generations (27). Since training on *P. vranovensis* resulted in an increase in *daf-7p::gfp* levels in the ASI (Figure 2C-D), and the adult F1 progeny of GRb0427-trained mothers showed robust avoidance of *P. vranovensis* compared to the F1 progeny of the control (OP50-trained) mothers (**Figure 1**) we examined the expression of *daf-7p::gfp* in progeny of GRb0427-trained mothers: these progeny express higher levels of *daf-7p::gfp* in the ASI neurons (**Figure 3A-B**) compared to that in F1 animals from OP50-trained mothers.

**Figure 3.**
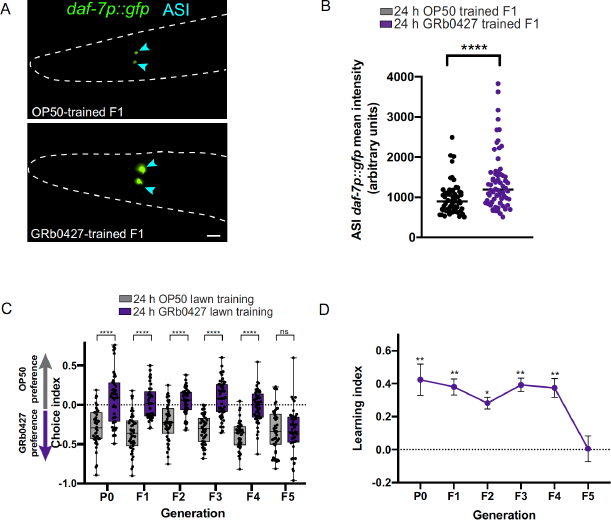
GRb0427-mediated learned avoidance is inherited transgenerationally. **(A)** F1 progeny of GRb0427-trained animals have higher *daf-7p::gfp* expression in the ASI sensory neurons (blue arrowheads). Scale bar = 10 μm. **(B)** Quantification of mean ASI *daf-7p::gfp* intensities from F1 progeny of OP50-trained and GRb0427 animals shows higher expression in F1 progeny of GRb0427-trained animals. **(C)** Untrained F1-F4 progeny of GRb0427-trained P0 animals avoid GRb0427 relative to the progeny of OP50-trained control P0 animals. This avoidance is lost in the F5 generation. **(D)** Learning index (naive choice index – trained choice index) of generations P0–F5. Error bars represent mean ± SEM. Each dot represents an individual choice assay plate (C) or an individual neuron for fluorescence images (B). Boxplots: center line, median; box range, 25th–75th percentiles; whiskers denote minimum-maximum values. Unpaired, two-tailed Student’s t test, ^****^p < 0.0001 (B); one-way ANOVA with Tukey’s multiple comparison’s test, ****p<0.0001, **p<0.01, *p<0.05, ns, not significant (C, D).

To examine if learned avoidance to GRb0427 is inherited transgenerationally (beyond the F1 generation), we tested the F2-F5 progeny for avoidance. The learned avoidance of GRb0427 lasts up to the F4 generation, but returns to naïve attraction to GRb0427 in the F5 generation (**Figure 3C-D**). Thus, GRb0427 training does induce transgenerational inheritance of learned avoidance behavior, as we previously found for PA14. Notably, in contrast to the higher avoidance of PA14 in the P0 generation than in F1-F4 (27), the level of avoidance of *P. vranovensis* is constant across P0 through F4 (Figure 3C). This result is consistent with the *daf-7p::gfp* and *irg-1p::gfp* results (Figure 2E-F) suggesting that innate immunity pathways may not contribute significantly to *C. elegans’* avoidance of *P. vranovensis*, but rather that the major pathway of avoidance in P0s is through the same pathway as in F1-F4.

### *P. vranovensis* small RNAs drive learned avoidance

Since learned avoidance to *P. vranovensis* is transgenerationally inherited, and trans-generational inheritance of avoidance of PA14 is driven by its small RNA, P11, we next asked if small RNAs made by *P. vranovensis* induce avoidance. Like PA14, when adult *C. elegans* were exposed for 24 hours to sRNAs isolated from *P. vranovensis* GRb0427, worms learned to avoid *P. vranovensis* (**Figure 4A**). Exposure to *P. vranovensis* sRNAs increased *daf-7p::gfp* expression in the ASI sensory neurons (**Figure 4B-C**). We next tested if *P. vranovensis* sRNA-induced learned avoidance is transgenerationally inherited. Indeed, as observed for *P. vranovensis* lawn exposure, *P. vranoven-sis* sRNA-induced avoidance is inherited up to the F4 generation and resets in the F5 (**Figure 4D-E**).

**Figure 4.**
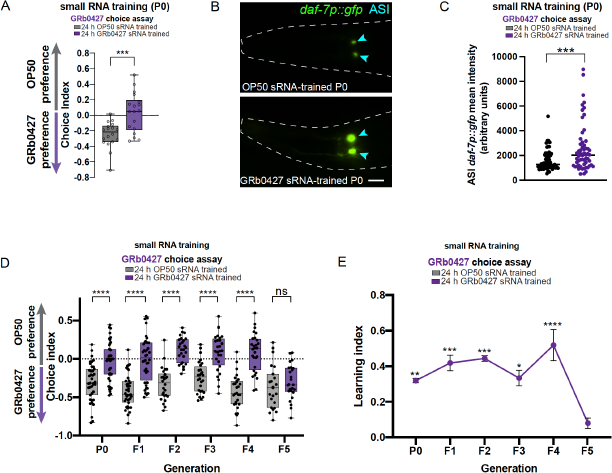
GRb0427 small RNAs induce learned avoidance and its transgenerational inheritance. **(A)** Worms trained on GRb0427 small RNAs exhibit learned avoidance of GRb0427 in an OP50-GRb0427 choice assay. **(B)** GRb0427 sRNA-trained animals have higher *daf-7p::gfp* expression in ASI sensory neurons (blue arrowheads). Scale bar = 10 μm. **(C)** Quantification of mean ASI *daf-7p::gfp* intensities from OP50 sRNA-trained and GRb0427 sRNA-trained animals shows higher expression in GRb0427 sRNA-trained animals. **(D)** Untrained F1-F4 progeny of GRb0427 sRNA-trained P0 animals avoid GRb0427 relative to the progeny of OP50 sRNA-trained control P0 animals. This avoidance is lost in the F5 generation. **(E)** Learning index (naive choice index – trained choice index) of generations P0–F5. Error bars represent mean ± SEM. Each dot represents an individual choice assay plate (A, D) or an individual neuron for fluorescence images (C). Boxplots: center line, median; box range, 25th–75th percentiles; whiskers denote minimum-maximum values. Unpaired, two-tailed Student’s t test (A, C), ***p<0.001; one-way ANOVA with Tukey’s multiple comparison’s test, ^****^p < 0.0001, ***p<0.001, **p<0.01, *p<0.05, ns, not significant (D, E).

### A specific *P. vranovensis* sRNA induces learned avoidance

We next asked if learned avoidance induced by *P. vranovensis* small RNAs is species-specific. We trained worms on *P. vranovensis* sRNAs and tested avoidance of PA14; *P. vranovensis* sRNA training induces avoidance to PA14, similar to training on PA14 sRNAs (**Figure 5A**). Intriguingly, worms exposed to bacteria expressing only the PA14 sRNA P11 also induces avoidance to *P. vranovensis* (**Figure 5B**), suggesting that the underlying mechanism in *P. vranovensis* sRNA-induced avoidance is like that of PA14-induced avoidance. Consistent with this idea, loss of *maco-1*, either by mutation (**Figure 5C**) or by RNAi treatment (**Figure 5D**) significantly reduces the strong naïve preference for GRb0427. Similarly, exposure to *P. vranovensis* small RNAs does not further increase *P. vranovensis* avoidance (**Figure 5C**). Quantitative RT-PCR showed a decrease in relative *maco-1* transcript abundance in GRb0427-trained animals (**Figure 5E**). This observation indicates that learned avoidance induced by *P. vranovensis* targets *maco-1*, as PA14’s P11 small RNA does. We also examined differentially-expressed genes between *P. vranovensis*-treated and *E. coli* HB101-treated P0 adult worms in RNA sequencing data reported in Burton et al., 2020 (8); consistent with our results, *maco-1* levels are reduced in *P. vranovensis*-treated animals (log_2_fold change = -0.2837998, padj = 0.027) in this independent analysis.

**Figure 5.**
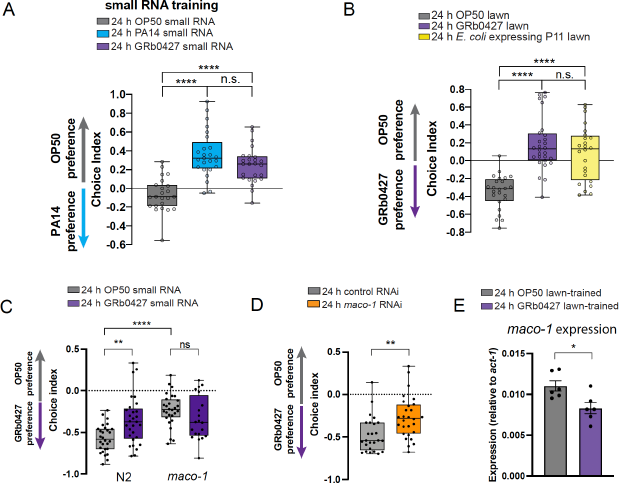
GRb0427 sRNA-induced avoidance requires *maco-1*. **(A)** Worms were trained on OP50 (gray), *Pseudomonas aeruginosa* PA14 (blue), or GRb0427 (purple) small RNAs, and tested for their PA14 preference in a bacterial choice assay between OP50 and PA14. Worms treated on GRb0427 small RNAs not only avoid GRb0427 (Figure 3A), but also avoid PA14 in an OP50-PA14 choice assay. **(B)** Worms trained on the PA14 small RNA P11 avoid GRb0427 in an OP50-GRb0427 choice assay. **(C)** N2 (wild type) worms trained on GRb0427 small RNAs avoid GRb0427. *maco-1(ok3165)* loss-of-function mutant worms naively avoid GRb0427 and do not show increased avoidance upon exposure to GRb0427 sRNAs. **(D)** Upon downregulation of *maco-1* by RNAi, wild type worms exhibit higher naïve avoidance of GRb0427 compared to control RNAi-treated wild-type worms. **(E)** Relative transcript abundance of *maco-1* (relative to *act-1* mRNA, used as the house-keeping gene for reference) in OP50 AND GRb0427-treated animals. Each data point represents an independent biological replicate, and 3 technical replicates were used for each biological replicate. Data are represented as mean +/- S.E.M. Each dot represents an individual choice assay plate (A-D). Boxplots: center line, median; box range, 25th–75th percentiles; whiskers denote minimum-maximum values, One-way ANOVA with Tukey’s multiple comparison’s test, ****p<0.0001, ns, not significant (A,B); Two-way ANOVA with Tukey’s multiple comparison’s test, **p<0.01, ****p<0.0001, ns, not significant (C); Unpaired, two-tailed Student’s t test, **p<0.01 (D), *p<0.05 (E).

We next examined whether *P. vranovensis* might encode a P11-like small RNA. We analyzed the recently-sequenced genome of *P. vranovensis* (8) and did not find any genomic region with sequence homology to P11. In fact, there is no region analogous to the operon that contains P11 in the *P. vranovensis* genome. That is, while P11 can induce similar avoidance of GRb0427 as it does for PA14, and GRb0427 induces avoidance through a small RNA, GRb0427 does not appear to encode a small RNA with a P11-like sequence in its genome; therefore, we needed to determine whether GRb0427 expresses a different sRNA that induces learned avoidance and inheritance of this avoidance.

While a P11-like sRNA cannot account for the learned avoidance of GRb0427, our small RNA *maco-1* experiments suggested that an sRNA with similarity to *maco-1* might be involved. Therefore, we searched the *P. vranovensis* genome for similarity to the *maco-1* sequence; we found five perfect matches to the *maco-1* coding region in the *P. vranovensis* genome, but only one of these, a 16nt match, lies in an intergenic region that would be likely to encode a small RNA (**Figure 6A, B**). Interestingly, this sequence identity lies in a different exon of *maco-1* (Exon 1) from P11’s 17nt perfect match (Exon 8). The *P. vranovensis* intergenic region containing this 16-nucleotide sequence match to *maco-1* is flanked by bacterial protein coding genes with predicted functions in the iron metabolism and sugar transport pathways (**Figure 6B**). We expressed a 347 bp region of this intergenic sequence (IntReg) in *E. coli*, and found that training on IntReg induces avoidance to *P. vranovensis* (**Figure 6C**). The avoidance induced by *E. coli* expressing the intergenic region persists for four generations after parental exposure (**Figure 6D**), and is lost by the F5 generation, similar to transgenerational inheritance of learned avoidance induced by a *P. vranovensis* lawn or small RNA exposure. We next investigated if the intergenic region encodes a small RNA. While the genome of *P. vranovensis* is published, no information on small RNAs were publicly available, so we sequenced the *P. vranovensis* total small RNA pool that induces learned avoidance and its transgenerational inheritance. Indeed, we detected a small RNA within the intergenic region that contains the 16-nucleotide sequence match to *maco-1* (**Figure 7A)**. This small RNA, which we named “Pv1”, is 124 bp long, and its homology to *maco-1*, like P11, lies in a predicted stem loop. We next asked whether exposure to Pv1 expressed in *E. coli* would be sufficient to induce avoidance to *P. vranovensis;* indeed, training on *E. coli-Pv1* induces avoidance in the mother generation (P0; **Figure 7B**), as well as the inter (F1)- and transgenerational (F2) inheritance of this learned avoidance (**Figure 7C**).

**Figure 6.**
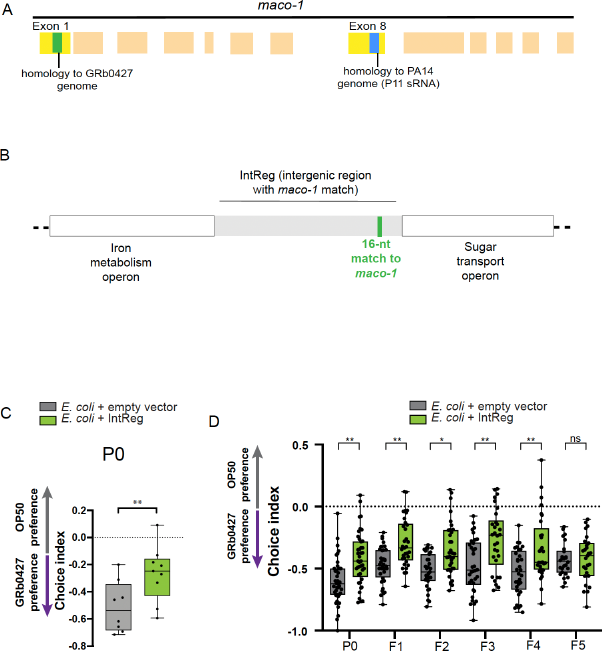
An intergenic region in the GRb0427 genome contains a 16-nucleotide perfect match to *maco-1* and is sufficient for learned avoidance of GRb0427. **(A)** The PA14 small RNA P11 contains a 17-nucleotide perfect match to the worm neuronal gene *maco-1* (Exon 8). By annotating the GRb0427 genome, we discovered an intergenic region with a 16-nucleotide perfect match to a stretch of Exon 1 of *maco-1*. **(B)** The GRb0427 genome has an intergenic region (flanked by an iron metabolism operon and a sugar transport operons) containing a 16-nucleotide perfect match to *maco-1*. **(C)** Training worms on *E. coli* expressing the intergenic region with the match to *maco-1* induces avoidance of GRb0427. **(D)** Untrained F1-F4 progeny of worms trained on *E. coli* expressing the intergenic region with the match to *maco-1* exhibit higher avoidance of GRb0427 compared to controls. This higher avoidance is lost in the F5 generation. Each dot represents an individual choice assay plate (C, D). Boxplots: center line, median; box range, 25th–75th percentiles; whiskers denote minimum-maximum values. Unpaired, two-tailed Student’s t test, ^**^p < 0.01, (C); one-way ANOVA with Tukey’s multiple comparisons test, ^**^p < 0.01, *p<0.05 ns, not significant (D).

**Figure 7.**
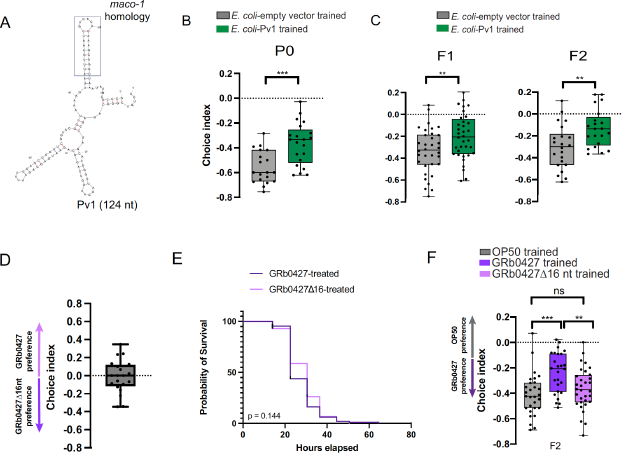
Pv1, a GRb0427 sRNA, is necessary and sufficient for transgenerational inheritance of learned avoidance. **(A)** mFold structure prediction for Pv1; the *maco-1* region is in a predicted stem loop of Pv1 (the boxed region). **(B)** Training worms on *E. coli* expressing just the Pv1 small RNA induces avoidance of GRb0427 (compared to training on *E. coli* expressing a control empty vector). **(C)** *E. coli*-Pv1 induced learned avoidance of GRb0427 is transgenerationally inherited. Untrained F1 and F2 progeny of *E. coli-*Pv1 trained P0 worms also exhibit higher GRb0427 avoidance compared to controls. **(D)** Naïve worms do not exhibit a preference towards either wild type GRb0427 or GRbΔ16nt strain in a choice assay. **(E)** Survival of worms on lawns of GRbΔ16nt and wild type GRb0427 are not significantly different. **(F)** F2 progeny of GRbΔ16nt-trained worms do not exhibit learned avoidance of GRb0427. Each dot represents an individual choice assay plate (C, D, F, H). Boxplots: center line, median; box range, 25th–75th percentiles; whiskers denote minimum-maximum values. Unpaired, two-tailed Student’s t test, ^**^p < 0.01, ***p<0.001, (B, C); One-way ANOVA with Tukey’s multiple comparison’s test, ^**^p < 0.01, ***p<0.001, ns, not significant (F). For the survival assay in (E), ns – not significant (by Log-rank (Mantel-Cox) test for survival).

To determine if Pv1’s sequence identity to *maco-1* is necessary for the learned avoidance to *P. vranovensis* and its transgenerational inheritance, we constructed a mutant *P. vranovensis* bacterial strain lacking the 16-nucleotide match to *maco-1*, Δ16, Although this Δ16 mutant GRb0427 strain is equally attractive to worms as wild-type GRb0427 (**Figure 7D**) and is similarly pathogenic to worms as wild-type GRb0427 **(Figure 7E)**, Δ16 is unable to induce transgenerational inheritance of learned avoidance (**Figure 7F)**. Together, our data suggest that a specific *P. vranovensis* small RNA, Pv1, is sufficient and its region of homology to *maco-1* is necessary for sRNA-mediated learned avoidance of GRb0427 and its transgenerational inheritance.

### *C. elegans* avoid the beneficial bacterium *Pseudomonas mendocina* following exposure to GRb0427 or Pv1

Transgenerational inheritance of learned avoidance of both PA14 and GRb0427 lasts for four generations, suggesting that there may be some benefit to “forgetting” learned avoidance of a pathogen. In the wild, worms experience a range of temperatures, and bacterial species can be differentially pathogenic at different temperatures. In the case of *P. aeruginosa* PA14, pathogenicity decreases at lower temperatures (23,59); thus, continuing to avoid this bacteria even at lower temperatures might cause the worms to mistakenly avoid a potentially good food source, and the PA14 small RNA, P11 is not expressed at lower temperatures (23). We wondered if this was also the case for GRb0427 and Pv1, or if other factors, including other members of the microbiome, might influence the cessation of avoidance.

We first asked if temperature affects the pathogenicity of GRb0427. Although less pathogenic to worms than 25°C-grown GRb0427, 15°C-grown GRb0427 still kills all the worms in the population before the control (OP50)-grown worms have started dying (**Figure 8A**). Unlike the differential expression of PA14’s P11 sRNA at different temperatures, the Pv1 small RNA is expressed in total RNA pools from both 25°C GRb0427 and 15°C GRb0427 (**Figure S2**).

**Figure 8.**
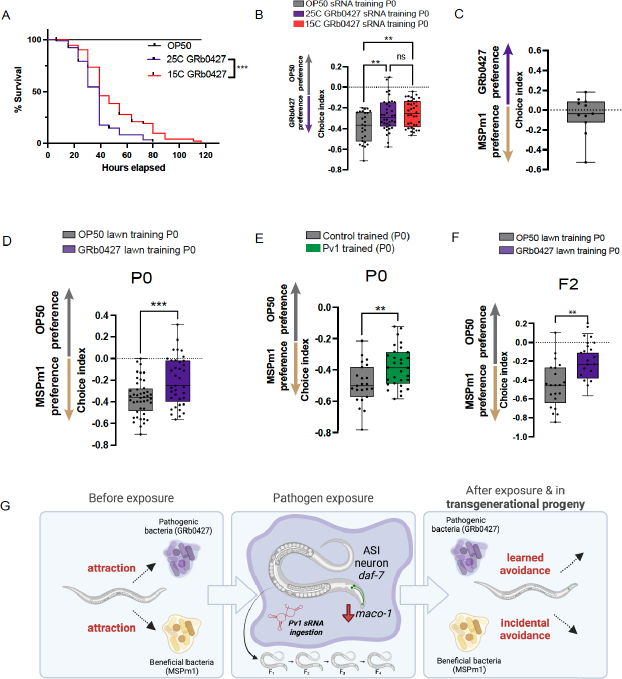
Pv1-induced learned avoidance causes avoidance of *P. mendocina*, a beneficial natural bacterial species. **(A)** Worms have significantly lower survival on a 25°C-grown GRb0427 lawn compared to a 15°C-grown GRb0427 lawn, but in both these conditions, all worms die before worms on an OP50 lawn have started dying, indicating that both 25°C and 15°C-grown GRb0427 are pathogenic. **(B)** Training worms on sRNAs from both 25°C and 15°C-grown GRb0427 induce learned avoidance. **(C)** Untrained worms have similar preference for GRb0427 and MSPm1 in a choice assay. **(D)** Training worms on a GRb0427 lawn reduces attraction to *P. mendocina* compared to OP50-trained control worms. **(E)** Training worms on *E. coli* expressing Pv1 reduces attraction to *P. mendocina*. **(F)** Naïve F2 progeny of GRb0427-trained P0 animals exhibit reduced attraction to *P. mendocina* relative to F2 progeny of OP50-trained P0 animals. **(G)** Schematic outlining the findings of the study. Prior to exposure to pathogenic GRb0427, worms are naively attracted to both GRb0427, and the non-pathogenic *P. mendocina*. Exposure to GRb0427 or to the Pv1 small RNA of GRb0427 induces learned avoidance of GRb0427 in worms and four generations of progeny. Exposed worms not only avoid the pathogenic GRb0427, but also the non-pathogenic *P. mendocina*. Each dot represents an individual choice assay plate (B-F). Boxplots: center line, median; box range, 25th–75th percentiles; whiskers denote minimum-maximum values. One-way ANOVA with Tukey’s multiple comparison’s test, ^**^p < 0.01, ns, not significant (B); Unpaired, two-tailed Student’s t test, ^**^p < 0.01, ***p<0.001 (D-F). For the survival assay in (A), ***p<0.001 (by Log-rank (Mantel-Cox) test for survival).

Consistent with Pv1 being expressed under both temperature conditions, worm populations trained on 25°C and 15°C GRb0427 small RNAs both avoid GRb0427 compared to worms trained on control (OP50) small RNAs (**Figure 8B**). Thus, it seems unlikely that temperature-dependent changes and Pv1 sRNA expression in GRb0427 pathogenicity drive the loss of memory of learned avoidance. We next asked if GRb0427-induced learned avoidance might alter worms’ responses to other bacteria. Several bacterial species in the worm microbiome are beneficial; that is, they increase worm growth rates and progeny production and extend lifespan (47). In fact, all the wild bacteria we tested are more attractive than OP50 (Figure 1). This preference likely evolved as *Pseudomonas* bacteria are sources of nutrition to *C. elegans* in the wild (47). Therefore, we asked if exposure to the pathogenic GRb0427 might cause worms to inadvertently avoid beneficial bacteria in their microbiome. Some of the beneficial bacterial species in the worm microbiome, e.g., *Pseudomonas mendocina*, belong to the same genus as GRb0427 (50); *C. elegans* are attracted to *Pseudomonas mendocina* (56). As we showed above, worms trained on a strain of *P. mendocina* from the worm’s microbiome, MSPm1, do not exhibit an altered response to *P. mendocina* compared to control (*E. coli* OP50)-trained worms (**Figure 1H**), as might be expected for beneficial bacteria. Consistent with this logic, we found that the *P. mendocina* genome does not encode the Pv1 small RNA used by GRb0427 to elicit avoidance, and the intergenic regions in the genome do not contain any significant contiguous matches to the Pv1 target *maco-1*. We next tested the effect of GRb0427 exposure on worms’ responses to *P. mendocina*. In their naïve state, *C. elegans* exhibits no preference for GRb0427 or MSPm1 over one another (**Figure 8C**). However, worms trained on GRb0427 for 24 hrs have significantly reduced attraction to *P. mendocina* compared to worms grown on the control *E. coli* OP50 (**Figure 8D**) – that is, although they have never been exposed to *P. mendocina*, and it is a beneficial food source they are naively attracted to (Figure 1H), exposure to pathogenic *P. vranovensis* induces avoidance of the beneficial bacteria. Similarly, training on *E. coli* expressing the Pv1 small RNA is sufficient to reduce attraction of worms to the beneficial *P. mendocina* (**Figure 8E**). Since Pv1 mediates transgenerational inheritance of learned GRb0427 avoidance, we next tested if altered *P. mendocina* preference upon GRb0427 training is inherited for multiple generations. Indeed, F2 progeny of GRb0427-trained animals continue to exhibit reduced attraction to *P. mendocina* **(Figure 8F)**.

Taken together, our results suggest that in the wild, Pv1 small RNA-mediated learned avoidance may protect worms and their progeny from the pathogenic GRb0427, but since this also results in avoidance of beneficial bacteria, it may be beneficial to reverse this memory after a few generations, when the worms may no longer encounter the original pathogen.

## Discussion

*C. elegans’* wild environments are rife with diverse bacterial species, and recent studies have provided comprehensive descriptions of *C. elegans’* natural microbial environments (43,44,47). The effects of these bacterial species on *C. elegans’* nervous system and behavior have been largely unexplored, and whether these bacteria can exert transgenerational effects on worms was previously unknown. Here we show that a small RNA from *Pseudomonas vranovensis* GRb0427, a pathogenic bacterial species found in *C. elegans’* natural environment, induces learned avoidance in worms. This learned avoidance persists for four generations, similar to the learned avoidance induced by the P11 small RNA expressed in the clinical isolate of *P. aeruginosa*, PA14. This wild bacterial species, *P. vranovensis*, however, does not express the P11 small RNA; instead, it uses a distinct small RNA, which we identified and named Pv1, to induce avoidance and its transgenerational inheritance. The Pv1 small RNA, like P11, regulates avoidance by targeting the neuronal gene *maco-1* – but in a different exon - and subsequently upregulating the expression of the TGF-beta ligand *daf-7*.

Our study provides an example of small RNA-mediated cross-kingdom signaling between bacteria and their animal hosts, adding to the diverse instances of RNA-based trans-kingdom signaling (23,60,61). The identification of Pv1 expands the repertoire of bacterial small RNAs that can affect host physiology and begins to identify common principles of bacterial small RNA-mediated regulation of *C. elegans* behavior. Both Pv1 and P11 target *maco-1*, a conserved ER-localized calcium regulator, indicating that the sRNA/*maco-1/daf-7* axis may be important for regulation by multiple bacterial small RNAs. Pv1, like P11, has a perfect match to its putative target, *maco-1*, indicating that bacterial small RNAs require perfect sequence complementarity for sRNA-mediated effects; moreover, these matches are present in stem loops of predicted secondary structures of these sRNAs (**Figure 6A**). However, Pv1 and P11 contain sequence matches to different exons of the *maco-1* gene sequence (**Figure 5A**), suggesting that the P11-matched region of *maco-1* is not the key element. Although P11 and Pv1 are similar in secondary structure and length, they have very low sequence homology, suggesting that these two small RNAs likely evolved independently. The odds of a *Pseudomonas* small RNA having a perfect 16 nt match to part of *maco-1* by chance are low (less than 0.03, which perhaps is an overestimate; see Methods). Therefore, although 16 nucleotide length is short, the perfect match requirement between a bacterial small RNA and a potential *C. elegans* transcript may sharply limit the possibilities. The fact that the *maco-1* sequence matches are present in predicted stem loops may also suggest that there may be convergent evolution towards a particular length or secondary structure. This supports the idea that several other bacterial sRNA-based regulation systems exist that may target the same or distinct *C. elegans* genes and may be yet to be discovered. Characterizing sRNAs from additional bacterial species that induce heritable behaviors in worms may help define molecular and structural features of sRNAs that can induce learning and inheritance, and may even allow us to predict which naturally existing exogenous sRNAs, including sRNAs from animal gut microbiomes, have the potential to induce heritable effects.

These findings establish bacterial small RNA-mediated regulation of host physiology as a phenomenon that may occur in the wild and in *C. elegans’* interaction with multiple different bacterial species. In fact, our data indicate that small RNAs may play a significant role in the neuronal response of *C. elegans* to wild bacteria: the behavioral response to GRb0427 is almost entirely mediated by small RNAs. In fact, the secondary metabolite-induced neuronal response observed upon *Pseudomonas aeruginosa* PA14 exposure (*daf-7* induction in ASJ neurons) (27,58) is not observed upon GRb0427 exposure, and the innate immunity gene *irg-1* (62) is not greatly induced, either, suggesting that the sRNA-mediated response may be the major mechanism of avoidance of this pathogen. The sRNA-mediated learned response is also mechanistically distinct from a previously-described intergenerational effect on progeny (embryo) survival upon *P. vranovensis* exposure, which does not appear to use bacterial small RNAs or continue beyond the F1 generation (8), or a P0 avoidance effect induced by *O. vermis* (63). Nor is there any overlap with the general response to pathogens (64), which is mediated by translational inhibition; for example, *vhp-1*, which plays a key role in the general response, has the opposite effect on pathogen avoidance (23). The transgenerational effect we observe is species-specific; in fact, of the pathogenic species we tested here (Figure 1), and in previous work (*S. marcescens* (27)), only PA14 and GRb0427, two *Pseudomonas* pathogens, induce transgenerational inheritance of pathogen avoidance. Although not every *Pseudomonas* species induces avoidance (the beneficial *P. mendocina* does not, for example), of the pathogens we have tested, only *Pseudomonas* pathogens seem to induce a small RNA-mediated transgenerational avoidance response. Further work will determine how broadly *C. elegans* employs this mechanism to avoid *Pseudomonas* pathogens.

Our results suggest that in the wild, Pv1 small RNA-mediated learned avoidance may protect worms and their progeny from the pathogenic GRb0427, but since this also results in avoidance of beneficial bacteria, it may be beneficial to reverse this memory after a few generations when the worms may no longer encounter the original pathogen. We showed that exposure to *P. vranovensis* GRb0427 alters *C. elegans’* response to multiple bacterial species; not only do they learn to avoid the pathogenic GRb0427, but they also (mistakenly) avoid the beneficial *P. mendocina*. In the wild, it might be beneficial to inherit this transgenerational memory for only a limited number of generations, rather than indefinitely, to prevent the deleterious avoidance of nutrient sources. After exposure to a pathogen such as GRb0427, learned avoidance memory might protect the next few progeny generations from the pathogen, but it might be beneficial to revert this memory, since the environmental microbial composition in the wild may subsequently change to include beneficial bacteria. Reversal of this memory may have evolved to protect worms from maladaptive avoidance of other beneficial bacterial species once the pathogen threat has passed. Since most bacterial families in *C. elegans’* microbiome include beneficial as well as detrimental members that may present similar odors to the worms, reversible transgenerational responses to multiple other bacterial species may have evolved as an adaptive strategy.

Together, our results suggest that the “interpretation” of bacterial small RNA signals may in fact be one reason that *C. elegans* developed such robust RNA interference mechanisms: in addition to endogenous RNAi systems used to silence errant germline transcripts (65–67), mechanisms to regulate and process exogenous sRNAs from the worm’s environment and bacterial food sources may be critical for avoidance of pathogens in *C. elegans’* environment. If *C. elegans* constantly surveys its bacterial environment, but also needs to cease avoidance after a few generations to prevent accidentally avoiding beneficial food sources, the bacterial small RNA/RNA interference pathway may provide an ideal adaptable, re-settable response mechanism. More examples will be necessary to determine whether the system is restricted to *Pseudomonad* species and small RNAs that target *maco-1*, as well as to better define the characteristics of key bacterial small RNAs.

Our work provides proof of principle that transgenerational inheritance of learned avoidance is likely to benefit *C. elegans* in the wild. Although bacterial small molecules have been widely implicated in bacteria-host interactions (68), the potential roles of bacterial small RNAs in regulating host physiology are largely understudied. Identifying different bacterial small RNAs that can interact with host organ systems will lead to a better understanding of this new dimension of bacteria-host interactions.

## Acknowledgements

We thank Buck Samuel for sharing the GRb0427 and Jub38 bacterial strains, the *C. elegans* microbiome resource CeMbio for the remaining bacterial strains, and the *C. elegans* Genetics Center for worm strains; W. Wang, J. Arly Volmar, and J. Miller (Genomics Core Facility, Princeton University); BioRender for model figure design software; the Murphy lab for discussion and feedback; and J. Ashraf and W. Keyes for assistance. C.T.M. is the Director of the Glenn Center for Aging Research at Princeton and the Lewis Sigler Institute for Integrative Genomics. This work was supported by a Pioneer Award to C.T.M. (NIGMS DP1GM119167), a Transformative R01 (1R01AT011963-01) Award to Z.G. & C.T.M., CDCP 75D30122C15113 to C.M., T32GM007388 (NIGMS) support of R.S.M., a Damon Runyon Fellowship (DRG-2481-22) to T.S., and an NSF predoctoral award to J.S.

## Author contributions

T.S., J.S., R.M., R.K., Z.G., and C.T.M. planned experiments; T.S., J.S., R.M., and R.K. performed experiments; T.S., J.S., R.M., and R.K. performed data analysis; T.S., J.S., R.K., and J.M. generated reagents, T.S. and C.T.M. wrote the manuscript (with input from the other authors); C.M., Z.G., and C.T.M. provided funding and supervised the project.

## Methods

### Resource availability

Further information and requests for resources and reagents should be directed to and will be fulfilled by Coleen T. Murphy.

### Materials availability

Bacterial and *C. elegans* strains generated in this study are available on request.

## Experimental model and subject details

### Bacterial strains

The GRb0427 strain was a generous gift from Dr. Buck Samuel’s lab. OP50 was obtained from the C.G.C. The PA14 strain was a gift from Prof. Zemer Gitai’s lab. Control (L4440) and *maco-1* RNAi were obtained from the Ahringer RNAi library, and the respective sequences were verified. *P. mendocina* MSPm1 is part of the CeMbio (50) collection and was obtained from CGC.

### Engineered bacterial strains

*E. coli* expressing the GRb0427 347nt intergenic region containing the 16-nucleotide match to *maco-1* was constructed by Gibson Assembly. The entire intergenic region was cloned out of GRb0427 and ligated into pZE27GFP using a double restriction digestion to open the plasmid and a single fragment Gibson assembly to ligate. This plasmid was then transformed into MG1655 *E. coli* using a standard transformation protocol. *E. coli* expressing the Pv1 small RNA was constructed in a similar method, using primers specific to the Pv1 region.

The deletion of the 16 nt region of Pv1 (that matches to *C. elegans maco-1*) from the GRb0427 genome was constructed by two-step allelic exchange using plasmid pEXG2. Briefly, ∼600 bp fragments directly upstream and downstream of Pv1 sequence were amplified from GRb0427 gDNA using primer pairs (Pv1-KO1, Pv1-KO2) and (Pv1-KO3, Pv1-KO4), respectively. Overlap-extension PCR, with primer pair (Pv1-KO1, Pv1-KO4), was used to fuse together the upstream and downstream fragments. The final fragment was cloned into pEXG2, which was PCR linearized with primer pair (pEGX2-Lin1, pEGX2-seq2). The pEXG2 plasmid was integrated into GRb0427 by conjugation from donor strain *E. coli* s17 and exconjugants were selected on gentamycin 30 μg/mL and irgasan 100 μg/mL. Mutants of interest were counterselected on 15% sucrose and the proper deletion was confirmed via PCR, using primers (Pv1-seq1, Pv1-seq2), and sequencing of PCR products.

Primer details:

Pv1-KO1 - CGCACCCGTGGAAATTAATTGCTTCAGTGAAGGGCGG

Pv1-KO2 - TTCAGCATGCTTGCGGCTCGAGCACAGAAAGCAGATTAAATATGCGC

Pv1-KO3 - CTCGAGCCGCAAGCATGCTGAATTTTAGCCGTACCGAACAAGC

Pv1-KO4 - CCGGAAGCATAAATGTAAGCGTCCTTGTCGGGGC

pEGX2-Lin1 – CTTTACATTTATGCTTCCGGCTCGTA

pEGX2-Lin2 - AATTAATTTCCACGGGTGCGCATG

Pv1-seq1 - CGCACCCGTGGAAATTAATTGCTTCAGTGAAGGGCGG

Pv1-seq2 – CCGGAAGCATAAATGTAAGCGTCCTTGTCGGGGC

### *C. legans* strains

The following strains were used in this paper and were obtained from the CGC: N2 (wild type), AU133: *agls17[Pmyo-2::mCherry + Pirg-1::gfp]*, FK181: *ksIs2 [Pdaf-7p::gfp + rol-6(su1006)]*, CQ759: *maco-1(ok3165)* 2X outcrossed, and JU1580 (wild type, natural *C. elegans* isolate).

### Cultivation of bacterial strains

*E. coli* OP50, *P. vranovensis* GRb0427, *P. vranovensis* GRb0427 Δ16 nt, *Raoultella sp*. Jub38, the CeMbio strains, and *P. aeruginosa* PA14 were grown in LB (10 g/l tryptone + 5 g/l yeast extract + 10 g/l NaCl in distilled water) in a shaker at 250 rpm. *E. coli* expressing the Pv1 sRNA were grown in LB supplemented with 50 μg/mL Kanamycin. RNAi bacteria were grown in LB supplemented with 100 μg/ml carbenicillin and 12.5 μg/ml tetracycline.

### General maintenance of *C. elegans* strains

Worm strains were maintained at 20°C on high-growth medium (HGM) plates (3 g/l NaCl, 20 g/l bacto-peptone, 30 g/l bacto-agar in distilled water, with 4 mL/L cholesterol (5 mg/mL in ethanol), 1 mL/L 1 M CaCl_2_, 1 mL/L 1 M MgSO_4_ and 25 mL/L 1 M KPO_4_ buffer (pH 6.0) added to molten agar after autoclaving) on *E. coli* OP50.

## Method details

### Training plate preparation

Training plates were prepared by pipetting 800 uL of bacteria onto NGM (3 g/L NaCl, 2.5 g/L Bacto-peptone, 17 g/L Bacto-agar in distilled water, with 1 mL/L cholesterol (5 mg/mL in ethanol), 1 mL/L 1M CaCl2, 1 mL/L 1M MgSO4, and 25 mL/L 1M potassium phosphate buffer (pH 6.0) added to molten agar after autoclaving) or HG plates. Pathogenic bacteria such as PA14 or GRb0427 were prepared on NGM plates to avoid overgrowth, while sRNA producing MG1655 were prepared on HG plates. For sRNA training, 200 μl of OP50 was spotted in the center of a 6-cm NGM plate. Plates were stored at 25°C for 48hrs. RNAi bacteria were prepared on HG plates (supplemented with 1 mL/L 1M IPTG, and 1 mL/L 100 mg/mL carbenicillin) and kept at room temperature for 48 hrs. Plates were then taken out of the incubator and allowed to cool to room temperature before moving animals onto them.

### Bacterial choice assay plate preparation

Overnight bacterial cultures were diluted in LB to an OD_600_ = 0.5, and 25 μl of each bacterial suspension was spotted onto one side of a 60-mm NGM plate to make bacterial choice assay plates. These plates were incubated for 2 days at 25 °C.

### Preparation of bacteria for small RNA isolation

GRb0427 and *E. coli* OP50 bacteria were cultured for 16 hours overnight. 1 ml of either bacterial culture was diluted to an OD = 0.5, plated on 100mm NGM plates and grown at 25 °C for 48 hours. For 15 °C GRb0427 small RNA isolation, overnight (37°C) cultures of GRb0427 bacteria were centrifuged for 10 min at 5,000*g*. The supernatant was removed, and the remaining pellet was resuspended in 5 ml of fresh LB. Washed bacteria were used to inoculate (1:500) fresh LB to grow at 15 °C for 36 hrs. 1 ml of this bacterial culture was then diluted to an OD = 0.5, plated on 100mm NGM plates and grown at 15 °C for 48 hours.

Bacterial lawns were collected from the surface of the plates using a cell scraper. 1 ml of M9 buffer was applied to the surface of the bacterial lawn, and the bacterial suspension obtained by scraping was transferred to a 15-ml conical tube. *E. coli* OP50 from 15 plates, or GRb0427 from 10 plates were pooled in each tube and pelleted at 4500*g* for 8 min. The supernatant was discarded, and the bacterial pellet was resuspended in 900 uL of Trizol LS for every 100 μl of bacterial pellet.

The pellet was resuspended by vortexing and the tubes containing the bacterial pellet were frozen at −80 °C.

### Bacterial small RNA isolation

To isolate small RNA, bacterial pellets-Trizol suspensions were first incubated at 65 °C for 10 min with occasional vortexing. Debris were pelleted at 4500*g* for 5 min. The supernatant was transferred to 1.5 mL tubes (1 mL in each tube) and 200uL chloroform was added. Samples were mixed by inverting and centrifuged at 12,000*g* at 4 °C for 10 min. The aqueous phase obtained was used as input for small RNA extraction using the mirVana miRNA isolation kit. Extraction was done as per the manufacturer’s instructions for small RNA (<200 nt) isolation. Purified small RNA was used immediately in aversive learning assays or for sequencing, or frozen at −80 °C until further use.

### Training *C. elegans* on bacterial lawns and small RNAs

Wild-type N2 animals were synchronized by bleaching and grown until larval stage 4 (L4) on standard HG plates seeded with OP50. At L4 stage, they were transferred to training plates.

After 48 hrs at 25°C the training plates were left on the bench top for 30 mins to allow them to reach room temperature. For sRNA training, 100 ug of sRNA was added to the OP50 spot on the sRNA training plates. For RNAi, 200 μL of 0.1M IPTG was spotted onto seeded RNAi plates and left to dry at room temperature before adding worms. Larval stage 4 (L4) worms were washed off plates using M9 and left to pellet on the bench top for 2-3 min. Then, 5 μl of worms were placed onto sRNA-spotted training plates, 10 μl onto OP50 plates, or 20 μl onto RNAi plates, *E. coli* expressing Pv1, GRb0427, or GRb0427Δ16 training plates. Worms were incubated on training plates at 20°C in separate containers for 24 hours. After 24 hours, worms were washed off plates using M9 and washed an additional 2-3 times to remove excess bacteria. Trained worms were tested in the aversive learning assay.

### Aversive learning assay

On the day of the assay, bacterial choice assay plates were left at room temperature for 1 h before use. To start the assay, 1 μl of 1 M sodium azide was spotted onto each respective bacteria spot to be used as a paralyzing agent during choice assay. Worms were then washed off training plates in M9, allowed to pellet by gravity, and washed 2-3 additional times in M9. Using a wide orifice pipet tip, 5 μl of worms were spotted at the bottom of the assay plate, midway between the bacterial lawns. The assay plates were incubated at room temperature for 1 h. After that, the number of worms on each bacterial spot were counted. Plating a large number of worms (>200) on choice assay plates was avoided, because the worms clump at bacterial spots making it difficult to distinguish individual worms during counting, and also because high densities of worms can alter behavior.

For experiments testing behavior of the F1 generation, day 1 worms from parental (P_0_) training were bleached and eggs were placed onto HG plates and left for 3 days at 20 °C. After 3 days, the F1 worms (Day 1 – 72 hours) were washed off HG plates with M9. Some of the pooled worms were subjected to the aversive learning assay, and the remaining worms were bleached to obtain eggs. The eggs were then placed onto HG plates, which were left at 20 °C. After 3 days the F2 progeny were tested, and the same steps were followed for subsequent progeny generations.

### Annotation of the GRb0427 genome

The GRb0427 genome was downloaded from Burton et al., 2020 (8) and run through an annotation pipeline in python. This pipeline searches for all open reading frames across the genome on both strands and makes a temporary gene list. It then filters genes by both size and overlap with other genes to give a final predicted gene list. Our pipeline predicted 4952 genes in the genome. Multiple genes were then selected and run through NCBI BLAST to confirm their identities.

After identifying the genes in the genome, we used Standalone BLAST, specifically, BLAST-nShort, to identify regions of homology between *maco-1* and the GRb0427 genome. This yielded 5 hits: 4 hits of 16 nucleotides and 1 of 20 nucleotides. Next, these hits were run through a python function to determine if they lie in an intergenic region, and this filtered our list down to a singular 16 nucleotide hit. This hit lies within a predicted 347nt intergenic region.

### Bacterial small RNA sequencing

Prior to sRNA sequencing, each sample of GRb0427 sRNA was tested for *elegans* behavior. The size distribution of sRNA samples was examined on a Bioanalyzer 2100 using RNA 6000 Pico chip (Agilent Technologies). The sRNA sequencing protocol was similar to the protocol used in (23). Briefly, around 300 ng of sRNA from each sample was first treated with RNA 5′ pyrophosphohydrolase (New England Biolabs) at 37 °C for 30 min, then converted to Illumina sequencing libraries using the PrepX RNA-seq library preparation protocol on the automated Apollo 324 NGS Library Prep System (Takara Bio). The treated RNA samples were ligated to two different adapters at each end, then reverse-transcribed to cDNA and amplified by PCR using different barcoded primers. The libraries were examined on Bioanalyzer DNA High Sensitivity chips (Agilent) for size distribution, quantified by Qubit fluorometer (Invitrogen), and then pooled at equal molar amount and sequenced on Illumina NovaSeq 6000 S Prime flowcell as single-end 122-nt reads. The pass-filter reads were used for further analysis.

### Bacterial small RNA sequencing data analysis

4 replicates of GRb0427 small RNA and 3 replicates of *E. coli* OP50 small RNA were sequenced. Reads were mapped to the sequenced GRb0427 genome (8) using RNA STAR (69). Default settings were used for the RNA STAR mapping. The resulting BAM files were then loaded into IGV genome browser (70) for analysis of the intergenic region containing the 16-nucleotide sequence match to *maco-1*. The peaks and read strands indicated that 3 small RNAs lie in the intergenic region of interest, and one spans the 16nt region of match to *maco-1*. Using the Sapphire promoter analysis software for *Pseudomonas* species (71), we found highly confident predicted promoters that were consistent with the *C*. sequencing data. Of the 3 small RNAs one contains the 16 nt homology to *maco-1*, and this small RNA was named Pv1. The boundaries of Pv1 were determined based on the depletion of mapped reads at the same genomic positions across all four GRb0427 small RNA sequence datasets.

### Determination of the operon context of Pv1

We examined if Pv1 lies in a GRb0427 operon. Using the Operon Mapper operon detection software (72), we found that while Pv1 is flanked by operons for iron metabolism and sugar transport, Pv1 itself is not part of an operon.

### Pv1 structure prediction

The small RNA that has homology to *maco-1* was identified by our sequencing data to be 124 nucleotides in length. Using the mFold sRNA secondary structure prediction tool on the UNAFold webserver (73), we found that this RNA contains a long stem loop structure with the sequence match to *maco-1* beginning in a stem and ending after the turn of a loop. Interestingly P11’s sequence match to *maco-1* has a similar secondary structure context.

### Imaging and image analysis

*daf-7p::gfp* images of OP50, PA14, and GRb0427 worms were taken on a Nikon Eclipse Ti microscope. Worms were prepared and treated as described in ‘Worm preparation for training’. Worms were mounted on 1% agar pads on glass slides and immobilized using 1 mM levamisole. *Z*-stack multi-channel (DIC and GFP) images of day-1 adult GFP-transgenic worms were acquired at 60X magnification. Maximum intensity projections of head neurons were built using Fiji. Quantification of mean fluorescent intensity was done using NIS-Elements software. Average pixel intensity was measured in each worm by drawing a Bezier outline of the neuron cell body for 2 ASI head neurons.

For *irg-1p::gfp* quantification, worms were prepared as described in ‘Worm preparation for training’ and imaged at 20× magnification on a Nikon A1 R confocal microscope. Image analysis was done with Fiji, where ROIs were drawn around each animal in the field of vision and mean intensity values for all regions of interest were recorded and plotted.

### Survival assay

Survival assay on GRb0427 and GRb0427Δ16 lawns:

GRb0427 and GRb0427Δ16 bacteria were grown in liquid culture overnight (37°C) and diluted 1:4 to an OD = 0.5. 750 μL of diluted GRb0427 or GRb0427Δ16 was spread to completely cover six 6-cm NGM plates for each bacterial genotype. Plates were incubated for 2 days at 25°C to allow bacterial growth. Plates were equilibrated to 20°C before adding worms (84 hours post-bleach) to plates. Survival assays were performed at 20°C. The assay plates were counted every 6-9 h. Every 48h, worms were moved onto new plates.

Survival assay on 25°C-grown and 15°C-grown GRb0427 lawns:

Preparation of 25°C-grown GRb0427 survival assay plates - Overnight (37°C) cultures of GRb0427 bacteria were diluted in LB to (OD_600_) = 0.5 and used to fully cover six 6-cm nematode growth medium (NGM) plates. The plates were incubated for 2 days at 25°C.

Preparation of 15°C-grown GRb0427 survival assay plates - Overnight (37°C) cultures of GRb0427 bacteria were centrifuged for 10 min at 5,000g. The supernatant was removed, and the remaining pellet was resuspended in 5 ml of fresh LB. Washed bacteria were used to inoculate (1:500) fresh LB to grow at 15 °C for 36 hrs. Cultures were diluted in LB to an OD_600_ = 0.5 and used to seed NGM plates. The plates were incubated at 15 °C for 2 days.

Assay procedure - Plates were equilibrated to 20°C before adding Day 1 (72 hours post-bleach) worms to plates. Survival assays were performed at 20°C. The assay plates were counted every 6-9 h. Every 24h, worms were moved onto new plates of 25°C-grown and 15°C-grown GRb0427 (prepared as described above).

### PCR detection of the Pv1 small RNA

Total RNA and small RNA were extracted from 25°C-grown and 15°C-grown GRb0427 using the mirVana miRNA isolation kit, and reverse-transcribed to DNA (using SuperScript III First Strand Synthesis System). A 62 bp region and an 88 bp region of Pv1 were identified using the following PCR primers:

Pv1(62bp)Fwd:

CTGTGACGATTACAAATTAAC

Pv1(62 bp)Rev:

GCCGTACCGAACAAG

Pv1(88 bp)Fwd:

GCCTAGCACTGGTTAG

Pv1(88bp)Rev:

CTGTGACGATTACAAATTAAC

### Quantification of *maco-1* gene expression by qPCR

Worms were trained for 24 hours on *E. coli* OP50 and GRb0427 (as described in the ‘Training *C. elegans* on bacterial lawns and small RNAs’ section). After training, the worms were collected in M9 and washed several times to remove excess bacteria. Worm pellets were crushed in liquid nitrogen and transferred to an appropriate volume of Trizol LS (100 μL of worm pellet in 900 μL of Trizol). Total RNA was extracted from sample-Trizol suspensions using chloroform extraction, isopropanol and ethanol precipitation, and cleanup using the RNeasy mini kit. cDNA was made from 1 ug of RNA using the Superscript III First-strand system for RT-PCR. The extracted cDNA was used as input for qPCR reactions using the Power SYBR green qPCR master mix and protocol and run on a Viia7 Real-time PCR system. The qPCR primers used are listed below:

*maco-1* Forward:

GTGTCACGACAATTGCC

*maco-1* Reverse:

CACATAGGTAGTGGCGAG

*act-1* Forward:

GGCCCAATCCAAGAGAGGTATC (74)

*act-1* Reverse:

CAACACGAAGCTCATTGTAGAAGG (74)

### Statistical analysis

Survival assays were assessed using Log-rank (Mantel-Cox) tests. For the comparison of choice indices between more than two genotypes, one-way ANOVA with Tukey’s multiple comparisons test was used. For comparisons of choice indices between genotypes and between conditions (naïve vs learned), two-way ANOVA with Tukey’s multiple comparisons test was used. Unpaired t tests were performed for comparisons between two groups. Experiments were repeated on separate days with separate populations, to confirm that results were reproducible. Prism 9 software was used for all statistical analyses.

### Perfect match calculation

There are approximately 500 small RNAs in *P. aeruginosa* (75) with an average length of 188 nt, constituting a total of ∼86,000 16-nt windows within these small RNAs. The length of the *maco-1* coding sequence is approximately 2700 nucleotides, and so contains ∼1,350 semi-independent 16-nt windows (allowing up to 90% overlap between neighboring 16-nt windows). The product of these two numbers is ∼116,100,000 pairs of potentially matching windows. Dividing by 4^16 possible 16-nt sequences yields an estimated probability of ∼0.027. It should be noted that this number is likely an overestimate, as it assumes that the 86,000 windows in the small RNAs are independent of one another, which is not the case.

## Supplementary Figures

**Figure S1.**
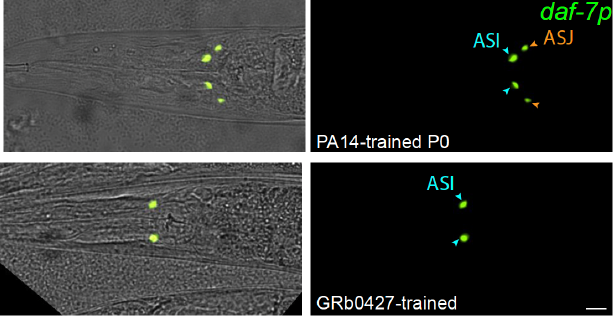
Representative image of a 24-hour PA14-trained (top) or GRb0427-trained (bottom) worm. *daf-7p::GFP* is expressed in the ASI (blue arrowheads) and ASJ (orange arrowheads) sensory neurons (right panel). Left panel is a merge of the GFP channel (right) and a brightfield channel. Scale bar = 10 μm.

**Figure S2.**
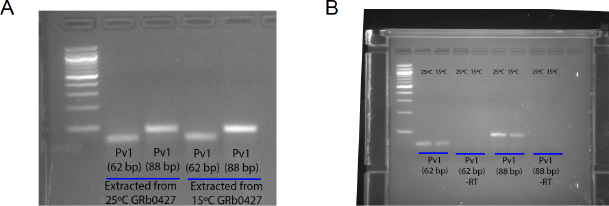
**(A)** 62 bp and 88 bp of Pv1 amplified by two different primer sets from total RNA pool extracted from 25°C and 15°C-grown GRb0427 (indicating expression of the Pv1 small RNA under both temperature conditions). **(B)** 62 bp and 88 bp of Pv1 amplified by the same primer sets as in (A) (also see Methods) from small RNA pool extracted from 25°C and 15°C-grown GRb0427. +RT and -RT indicate presence and absence respectively of the reverse transcriptase enzyme.

## References

1. Brzezinka K, Altmann S, Czesnick H, Nicolas P, Gorka M, Benke E, et al. Arabidopsis FORGETTER1 mediates stress-induced chromatin memory through nucleosome remodeling. Weigel D, editor. eLife. 2016 Sep 28;5:e17061.

2. Burton NO, Greer EL. Multigenerational epigenetic inheritance: Transmitting information across generations. Semin Cell Dev Biol. 2022 Jul;127:121–32.

3. Cecere G. Small RNAs in epigenetic inheritance: from mechanisms to trait transmission. Febs Lett. 2021 Dec;595(24):2953–77.

4. Liberman N, Wang SY, Greer EL. Transgenerational Epigenetic Inheritance: From Phenomena to Molecular Mechanisms. Curr Opin Neurobiol. 2019 Dec;59:189–206.

5. Lim JP, Brunet A. Bridging the transgenerational gap with epigenetic memory. Trends Genet. 2013 Mar 1;29(3):176–86.

6. Sun H, Damez-Werno DM, Scobie KN, Shao NY, Dias C, Rabkin J, et al. ACF chromatin-remodeling complex mediates stress-induced depressive-like behavior. Nat Med. 2015 Oct;21(10):1146–53.

7. Burton NO, Willis A, Fisher K, Braukmann F, Price J, Stevens L, et al. Intergenerational adaptations to stress are evolutionarily conserved, stress-specific, and have deleterious trade-offs. eLife. 2021 Oct 8;10:e73425.

8. Burton NO, Riccio C, Dallaire A, Price J, Jenkins B, Koulman A, et al. Cysteine synthases CYSL-1 and CYSL-2 mediate C. elegans heritable adaptation to P. vranovensis infection. Nat Commun. 2020 Apr 8;11(1):1741.

9. Burton NO, Furuta T, Webster AK, Kaplan REW, Baugh LR, Arur S, et al. Insulin-like signalling to the maternal germline controls progeny response to osmotic stress. Nat Cell Biol. 2017 Mar;19(3):252–7.

10. Conine CC, Sun F, Song L, Rivera-Pérez JA, Rando OJ. Small RNAs Gained during Epididymal Transit of Sperm Are Essential for Embryonic Development in Mice. Dev Cell. 2018 Aug 20;46(4):470–480.e3.

11. Conine CC, Moresco JJ, Gu W, Shirayama M, Conte D, Yates JR, et al. Argonautes promote male fertility and provide a paternal memory of germline gene expression in C. elegans. Cell. 2013 Dec 19;155(7):1532–44.

12. Guida MC, Birse RT, Dall’Agnese A, Toto PC, Diop SB, Mai A, et al. Intergenerational inheritance of high fat diet-induced cardiac lipotoxicity in Drosophila. Nat Commun. 2019 Jan 14;10(1):193.

13. Hibshman JD, Hung A, Baugh LR. Maternal Diet and Insulin-Like Signaling Control Intergenerational Plasticity of Progeny Size and Starvation Resistance. PLOS Genet. 2016 Oct 26;12(10):e1006396.

14. Hong C, Lalsiamthara J, Ren J, Sang Y, Aballay A. Microbial colonization induces histone acetylation critical for inherited gut-germline-neural signaling. PLoS Biol. 2021 Mar;19(3):e3001169.

15. Jordan JM, Hibshman JD, Webster AK, Kaplan REW, Leinroth A, Guzman R, et al. Insulin/IGF Signaling and Vitellogenin Provisioning Mediate Intergenerational Adaptation to Nutrient Stress. Curr Biol CB. 2019 Jul 22;29(14):2380–2388.e5.

16. Lim AI, McFadden T, Link VM, Han SJ, Karlsson RM, Stacy A, et al. Prenatal maternal infection promotes tissue-specific immunity and inflammation in offspring. Science. 2021 Aug 27;373(6558):eabf3002.

17. Ow MC, Nichitean AM, Hall SE. Somatic aging pathways regulate reproductive plasticity in Caenorhabditis elegans. eLife. 2021 Jul 8;10:e61459.

18. Pereira AG, Gracida X, Kagias K, Zhang Y. C. elegans aversive olfactory learning generates diverse intergenerational effects. J Neurogenet. 2020;34(3–4):378–88.

19. Perez MF, Shamalnasab M, Mata-Cabana A, Valle SD, Olmedo M, Francesconi M, et al. Neuronal perception of the social environment generates an inherited memory that controls the development and generation time of C. elegans. Curr Biol. 2021 Oct 11;31(19):4256–4268.e7.

20. Willis AR, Zhao W, Sukhdeo R, Wadi L, El Jarkass HT, Claycomb JM, et al. A parental transcriptional response to microsporidia infection induces inherited immunity in offspring. Sci Adv. 2021 May;7(19):eabf3114.

21. Gammon DB, Ishidate T, Li L, Gu W, Silverman N, Mello CC. The Antiviral RNA Interference Response Provides Resistance to Lethal Arbovirus Infection and Vertical Transmission in Caenorhabditis elegans. Curr Biol CB. 2017 Mar 20;27(6):795–806.

22. Jobson MA, Jordan JM, Sandrof MA, Hibshman JD, Lennox AL, Baugh LR. Transgenerational Effects of Early Life Starvation on Growth, Reproduction, and Stress Resistance in Caenorhabditis elegans. Genetics. 2015 Sep;201(1):201–12.

23. Kaletsky R, Moore RS, Vrla GD, Parsons LR, Gitai Z, Murphy CT. C. elegans interprets bacterial non-coding RNAs to learn pathogenic avoidance. Nature. 2020 Oct;586(7829):445–51.

24. Legüe M, Caneo M, Aguila B, Pollak B, Calixto A. Interspecies effectors of a transgenerational memory of bacterial infection in Caenorhabditis elegans. iScience. 2022 Jul 15;25(7):104627.

25. Mondotte JA, Gausson V, Frangeul L, Suzuki Y, Vazeille M, Mongelli V, et al. Evidence For Long-Lasting Transgenerational Antiviral Immunity in Insects. Cell Rep. 2020 Dec 15;33(11):108506.

26. Moore RS, Kaletsky R, Lesnik C, Cota V, Blackman E, Parsons LR, et al. The role of the Cer1 transposon in horizontal transfer of transgenerational memory. Cell. 2021 Sep 2;184(18):4697–4712.e18.

27. Moore RS, Kaletsky R, Murphy CT. Piwi/PRG-1 Argonaute and TGF-β Mediate Transgenerational Learned Pathogenic Avoidance. Cell. 2019 Jun 13;177(7):1827–1841.e12.

28. Rechavi O, Houri-Ze’evi L, Anava S, Goh WSS, Kerk SY, Hannon GJ, et al. Starvation-Induced Transgenerational Inheritance of Small RNAs in C. elegans. Cell. 2014 Jul 17;158(2):277–87.

29. Rechavi O, Minevich G, Hobert O. Transgenerational inheritance of an acquired small RNA-based antiviral response in C. elegans. Cell. 2011 Dec 9;147(6):1248–56.

30. Toker IA, Lev I, Mor Y, Gurevich Y, Fisher D, Houri-Zeevi L, et al. Transgenerational inheritance of sexual attractiveness via small RNAs enhances evolvability in C. elegans. Dev Cell. 2022 Feb 7;57(3):298–309.e9.

31. Vogt MC, Hobert O. Starvation-induced changes in somatic insulin/IGF-1R signaling drive metabolic programming across generations. Sci Adv. 2023 Apr 7;9(14):eade1817.

32. Wang SY, Kim K, O’Brown ZK, Levan A, Dodson AE, Kennedy SG, et al. Hypoxia induces transgenerational epigenetic inheritance of small RNAs. Cell Rep. 2022 Dec 13;41(11):111800.

33. Webster AK, Jordan JM, Hibshman JD, Chitrakar R, Baugh LR. Transgenerational Effects of Extended Dauer Diapause on Starvation Survival and Gene Expression Plasticity in Caenorhabditis elegans. Genetics. 2018 Sep;210(1):263–74.

34. Baugh LR, Day T. Nongenetic inheritance and multigenerational plasticity in the nematode C. elegans. Wittkopp PJ, editor. eLife. 2020 Aug 25;9:e58498.

35. Sengupta T, Kaletsky R, Murphy CT. The Logic of Transgenerational Inheritance: Timescales of Adaptation. Annu Rev Cell Dev Biol. 2023;39(1):null.

36. Uller T, English S, Pen I. When is incomplete epigenetic resetting in germ cells favoured by natural selection? Proc R Soc B Biol Sci. 2015 Jul 22;282(1811):20150682.

37. Perez MF, Lehner B. Intergenerational and transgenerational epigenetic inheritance in animals. Nat Cell Biol. 2019 Feb;21(2):143–51.

38. Quadrana L, Colot V. Plant Transgenerational Epigenetics. Annu Rev Genet. 2016;50(1):467–91.

39. Santilli F, Boskovic A. Mechanisms of transgenerational epigenetic inheritance: lessons from animal model organisms. Curr Opin Genet Dev. 2023 Apr 1;79:102024.

40. Irazoqui JE, Troemel ER, Feinbaum RL, Luhachack LG, Cezairliyan BO, Ausubel FM. Distinct pathogenesis and host responses during infection of C. elegans by P. aeruginosa and S. taureus. PLoS Pathog. 2010 Jul 1;6(7):e1000982.

41. Pérez-Carrascal OM, Choi R, Massot M, Pees B, Narayan V, Shapira M. Host Preference of Beneficial Commensals in a Microbially-Diverse Environment. Front Cell Infect Microbiol. 2022;12:795343.

42. Stuhr NL, Curran SP. Bacterial diets differentially alter lifespan and healthspan trajectories in C. elegans. Commun Biol. 2020 Nov 6;3(1):653.

43. Berg M, Stenuit B, Ho J, Wang A, Parke C, Knight M, et al. Assembly of the Caenorhabditis elegans gut microbiota from diverse soil microbial environments. ISME J. 2016 Aug;10(8):1998–2009.

44. Dirksen P, Marsh SA, Braker I, Heitland N, Wagner S, Nakad R, et al. The native microbiome of the nematode Caenorhabditis elegans: gateway to a new host-microbiome model. BMC Biol. 2016 May 9;14:38.

45. Morgan E, Longares JF, Félix MA, Luallen RJ. Selective Cleaning of Wild Caenorhabditis Nematodes to Enrich for Intestinal Microbiome Bacteria. J Vis Exp JoVE. 2021 Aug 13;(174).

46. Petersen C, Dierking K, Johnke J, Schulenburg H. Isolation and Characterization of the Natural Microbiota of the Model Nematode Caenorhabditis elegans. J Vis Exp JoVE. 2022 Aug 17;(186).

47. Samuel BS, Rowedder H, Braendle C, Félix MA, Ruvkun G. Caenorhabditis elegans responses to bacteria from its natural habitats. Proc Natl Acad Sci. 2016 Jul 5;113(27):E3941–9.

48. Yang W, Petersen C, Pees B, Zimmermann J, Waschina S, Dirksen P, et al. The Inducible Response of the Nematode Caenorhabditis elegans to Members of Its Natural Microbiota Across Development and Adult Life. Front Microbiol [Internet]. 2019 [cited 2023 Jul 8];10. Available from: https://www.frontiersin.org/articles/10.3389/fmicb.2019.01793

49. Zhang F, Weckhorst JL, Assié A, Hosea C, Ayoub CA, Khodakova AS, et al. Natural genetic variation drives microbiome selection in the Caenorhabditis elegans gut. Curr Biol. 2021 Jun 21;31(12):2603–2618.e9.

50. Dirksen P, Assié A, Zimmermann J, Zhang F, Tietje AM, Marsh SA, et al. CeMbio - The Caenorhabditis elegans Microbiome Resource. G3 Bethesda Md. 2020 Sep 2;10(9):3025–39.

51. Zhang F, Berg M, Dierking K, Félix MA, Shapira M, Samuel BS, et al. Caenorhabditis elegans as a Model for Microbiome Research. Front Microbiol. 2017;8.

52. Lise Frézal, Marie Saglio, Gaotian Zhang, Luke Noble, Aurélien Richaud, Marie-Anne Félix. Genome-wide association and environmental suppression of the mortal germline phenotype of wild <em>C. elegans</em>. bioRxiv. 2023 Jan 1;2023.05.17.540956.

53. Haçariz O, Viau C, Karimian F, Xia J. The symbiotic relationship between Caenorhabditis elegans and members of its microbiome contributes to worm fitness and lifespan extension. BMC Genomics. 2021 May 19;22(1):364.

54. Kissoyan KAB, Drechsler M, Stange EL, Zimmermann J, Kaleta C, Bode HB, et al. Natural C. elegans Microbiota Protects against Infection via Production of a Cyclic Lipopeptide of the Viscosin Group. Curr Biol CB. 2019 Mar 18;29(6):1030–1037.e5.

55. Kissoyan KAB, Peters L, Giez C, Michels J, Pees B, Hamerich IK, et al. Exploring Effects of C. elegans Protective Natural Microbiota on Host Physiology. Front Cell Infect Microbiol [Internet]. 2022 [cited 2023 Jul 8];12. Available from: https://www.frontiersin.org/articles/10.3389/fcimb.2022.775728

56. Chen AJ, Zuazo C, Mellman K, Chandra R, L’Etoile N. C. elegans show Preference for Pseudomonas mendocina (MSPm1) and Proteus mirabilis (P. mirabilis sp?), and Repulsion to Pseudomonas lurida (MYb11); Growth on Pseudomonas mendocina (MSPm1) Increases Attraction to 2-nonanone. MicroPublication Biol [Internet]. 2022 Mar 4 [cited 2023 Jul 8]; Available from: https://www.micropublication.org/journals/biology/micropub-biology-000535

57. O’Donnell MP, Fox BW, Chao PH, Schroeder FC, Sengupta P. A neurotransmitter produced by gut bacteria modulates host sensory behaviour. Nature. 2020 Jul;583(7816):415–20.

58. Meisel JD, Panda O, Mahanti P, Schroeder FC, Kim DH. Chemosensation of Bacterial Secondary Metabolites Modulates Neuroendocrine Signaling and Behavior of C. elegans. Cell. 2014 Oct 9;159(2):267–80.

59. Tan MW, Mahajan-Miklos S, Ausubel FM. Killing of Caenorhabditis elegans by Pseudomonas aeruginosa used to model mammalian bacterial pathogenesis. Proc Natl Acad Sci. 1999 Jan 19;96(2):715–20.

60. Chow FWN, Koutsovoulos G, Ovando-Vázquez C, Neophytou K, Bermúdez-Barrientos JR, Laetsch DR, et al. Secretion of an Argonaute protein by a parasitic nematode and the evolution of its siRNA guides. Nucleic Acids Res. 2019 Apr 23;47(7):3594–606.

61. Buck AH, Coakley G, Simbari F, McSorley HJ, Quintana JF, Le Bihan T, et al. Erratum: Exosomes secreted by nematode parasites transfer small RNAs to mammalian cells and modulate innate immunity. Nat Commun. 2015 Oct 22;6:8772.

62. Estes KA, Dunbar TL, Powell JR, Ausubel FM, Troemel ER. bZIP transcription factor zip-2 mediates an early response to Pseudomonas aeruginosa infection in Caenorhabditis elegans. Proc Natl Acad Sci U S A. 2010 Feb 2;107(5):2153–8.

63. Petersen C, Pees B, Martínez Christophersen C, Leippe M. Preconditioning With Natural Microbiota Strain Ochrobactrum vermis MYb71 Influences Caenorhabditis elegans Behavior. Front Cell Infect Microbiol. 2021;11:775634.

64. Melo JA, Ruvkun G. Inactivation of conserved C. elegans genes engages pathogen- and xenobiotic-associated defenses. Cell. 2012 Apr 13;149(2):452–66.

65. Ketting RF, Haverkamp TH, van Luenen HG, Plasterk RH. Mut-7 of C. elegans, required for transposon silencing and RNA interference, is a homolog of Werner syndrome helicase and RNaseD. Cell. 1999 Oct 15;99(2):133–41.

66. Tabara H, Sarkissian M, Kelly WG, Fleenor J, Grishok A, Timmons L, et al. The rde-1 gene, RNA interference, and transposon silencing in C. elegans. Cell. 1999 Oct 15;99(2):123–32.

67. Vastenhouw NL, Fischer SEJ, Robert VJP, Thijssen KL, Fraser AG, Kamath RS, et al. A genome-wide screen identifies 27 genes involved in transposon silencing in C. elegans. Curr Biol CB. 2003 Aug 5;13(15):1311–6.

68. Bishai JD, Palm NW. Small Molecule Metabolites at the Host–Microbiota Interface. J Immunol. 2021 Oct 1;207(7):1725–33.

69. Dobin A, Davis CA, Schlesinger F, Drenkow J, Zaleski C, Jha S, et al. STAR: ultrafast universal RNA-seq aligner. Bioinformatics. 2013 Jan;29(1):15–21.

70. Thorvaldsdóttir H, Robinson JT, Mesirov JP. Integrative Genomics Viewer (IGV): high-performance genomics data visualization and exploration. Brief Bioinform. 2013 Mar 1;14(2):178–92.

71. Coppens L, Lavigne R. SAPPHIRE: a neural network based classifier for σ70 promoter prediction in Pseudomonas. BMC Bioinformatics. 2020 Sep 22;21(1):415.

72. Taboada B, Estrada K, Ciria R, Merino E. Operon-mapper: a web server for precise operon identification in bacterial and archaeal genomes. Bioinformatics. 2018 Dec 1;34(23):4118–20.

73. Zuker M. Mfold web server for nucleic acid folding and hybridization prediction. Nucleic Acids Res. 2003 Jul 1;31(13):3406–15.

74. Merritt C, Seydoux G. The Puf RNA-binding proteins FBF-1 and FBF-2 inhibit the expression of synaptonemal complex proteins in germline stem cells. Dev Camb Engl. 2010 Jun;137(11):1787–98.

75. Gómez-Lozano M, Marvig RL, Molin S, Long KS. Genome-wide identification of novel small RNAs in Pseudomonas aeruginosa. Environ Microbiol. 2012;14(8):2006–16.

